# *Staphylococcus aureus* planktonic but not biofilm environment induces an IFN-β macrophage immune response via the STING/IRF3 pathway

**DOI:** 10.1101/2023.05.30.542830

**Authors:** Elisabeth Seebach, Gabriele Sonnenmoser, Katharina F. Kubatzky

## Abstract

Chronic implant-related bone infections are a severe complication in orthopedic surgery. Biofilm formation on the implant surface impairs an effective immune response leading to bacterial persistence. In a previous study, we found that *Staphylococcus aureus* (SA) induced IRF3 activation and *Ifnb* gene expression only in its planktonic form but not in the biofilm. The aim of this study was to clarify the role of the stimulator of interferon genes (STING) in this process.

We treated murine RAW 264.7 macrophages with conditioned media (CM) generated from planktonic or biofilm cultured SA in combination with agonists or inhibitors of the cGAS/STING pathway. We further evaluated bacterial gene expression of planktonic and biofilm SA to find potential mediators. STING inhibition resulted in a loss of IRF3 activation and *Ifnb* induction in SA planktonic CM, whereas STING activation induced an IRF3 dependent IFN-β response in SA biofilm CM. Expression levels of genes associated with virulence decreased with biofilm formation while those associated with cyclic dinucleotide (CDN) synthesis did not correlate with *Ifnb* induction. We further observed that cGAS contributed to the *Ifnb* induction by SA planktonic CM although cGAS activation was not sufficient to induce *Ifnb* gene expression in SA biofilm CM.

Our data indicate that the different degrees of virulence associated with planktonic and biofilm SA environments result in an altered induction of an IRF3 mediated IFN-β response via the STING pathway. This finding suggests that the STING/IRF3/IFN-β axis is a potential candidate for further investigation as immunotherapeutic target in implant-related bone infections.

## Introduction

Chronicity of implant-related bone infections is associated with the formation of biofilms on the implant surface. Herein, the bacteria which are mainly of a staphylococci origin are protected against most antibiotics and host defense mechanisms [1, 2]. Additionally, the biofilm environment is discussed to shift the immune response towards tolerance further promoting bacterial persistence [2, 3]. This can either be caused by the increased metabolic activity of the biofilm bacteria leading to glucose deprivation and lactate accumulation in the local environment or by an adapted gene expression activity of the biofilm bacteria which results in a reduced production of immunogenic and toxic factors [4, 5].

In a previous study, we found the presence of bacterial factors to be more important for the different macrophage activation in a planktonic or biofilm environment than the respective glucose and lactate levels [6]. Additionally, we found that the stimulation with a *Staphylococcus aureus* (SA) planktonic environment led to the activation of the interferon regulatory factor 3 (IRF3) and *Ifnb* induction which was absent in the respective biofilm environment. Toll-like receptor (TLR)-2 and -9 activation was not the cause for the observed IRF3 phosphorylation and the underlying mechanism remained unclear. IRF3 can be also get activated through the cytosolic cyclic dinucleotide (CDN) sensor molecule STING (stimulator of interferon genes) [7, 8]. The CDNs c-di-AMP, c-di-GMP and 3’3’cGAMP are of bacterial origin and can directly interact with STING [9, 10]. These molecules serve as secondary messenger and are involved in bacterial homeostasis and environmental adaption, biofilm formation and virulence [11]. The eukaryotic CDN 2’3’-cGAMP is produced by the cyclic GMP-AMP synthase (cGAS) in response to cytosolic DNA and acts as an endogenous ligand for STING [12, 13]. The cytosolic DNA can either be of a microbial origin or released from the mitochondria or the nucleus upon cell damage [14]. Downstream of STING, TANK- binding kinase 1 (TBK1)-dependent phosphorylation leads to the activation of IRF3 and subsequent induction of type I interferon gene expression [15]. Furthermore, STING can activate the canonical NF-κB pathway thereby inducing inflammation-associated gene expression [8, 16]. The STING pathway is investigated in diverse chronic diseases such as cancer, autoimmune disorders as well as bacterial infections [17] and is of high therapeutic interest as a potential target for immunomodulatory approaches [18, 19]. Also, a contribution of STING in host defense mechanisms against SA infections have been described [20]. Deletion of STING showed opposite effects on the bacterial control between the different infection models [21–23]. Additionally, a direct activation of STING by c-di-AMP released from biofilm SA and subsequent IFN-β production was shown which was connected to anti- inflammatory macrophage polarization and bacterial persistence [24]. Nevertheless, the role of STING in the adaption of the immune response during the transition from an acute to chronic infection associated with biofilm formation is unclear and a direct comparison of the impact of STING activation on the immune response against planktonic or biofilm SA is missing.

The aim of our study was to investigate if STING plays a role in the IRF3 mediated induction of *Ifnb* gene expression in a SA planktonic environment and why this response cannot be detected in the respective biofilm environment. Therefore, we stimulated RAW 264.7 cells with conditioned media (CM) from SA planktonic and biofilm cultures in the presence of the STING inhibitor H-151 or STING ligand 3’3’-cGAMP, respectively and evaluated IRF3 phosphorylation, *Ifnb* induction and downstream IFN-β pathway activation. We further compared bacterial gene expression levels of CDN synthases, quorum sensing (QS) molecules and virulence associated factors between planktonic and biofilm cultured SA. Additionally, we investigated a contribution of cGAS to the *Ifnb* induction and if the difference in *Ifnb* induction between planktonic or biofilm CM was conserved between different SA strains.

## Materials and Methods

### Bacteria culture and preparation of conditioned media

*Staphylococcus aureus* strain ATCC 49230 (UAMS-1, isolated from a patient with chronic osteomyelitis) [25] was used for preparation of conditioned media (CM) as already described in a previous study [6]. SA was cultured on Columbia agar plates with 5% sheep blood (BD, Germany). On day of CM setup, 3 to 5 colonies were transferred into trypticase soy bouillon (TSB; BD, Germany) and cultivated under shaking for 3 hours at 37 °C to receive growth state bacteria. 6*10^5^ CFU/ml were adjusted in Dulbecco’s Modified Eagle Medium (DMEM) high glucose (Anprotec, Germany) + 10% heat-inactivated fetal calf serum (FCS; PAN-Biotech, Germany). For planktonic culture, bacteria were cultivated under shaking (200 rpm) for 24 hours at 37°C and 5% CO_2_. For biofilm culture, bacteria were plated in 24 well with 1 ml of cell culture medium with 10% FCS per well and cultivated under static conditions for 6 days with daily medium exchange. For CM, planktonic medium after 24 hours of culture or the last 24 hours medium change before day 6 biofilm culture was harvested by centrifugation at 4000 rpm for 15 min at 4°C. Harvested media were streaked onto agar plates and cultivated over night to ensure no contamination by other bacteria. CM were sterile filtered through a 0.2 µm filter, pH adjusted to physiological pH of growth medium and stored in aliquots at -80°C. For the unstimulated CM control, the growth medium (DMEM high glucose + 10% FCS) of the respective approach was treated similar to CM but without bacteria inoculation. For SA strain comparison, USA300 FPR3757, ATCC 25923 and SA113 were included. CM of these strains were prepared as described above. Here, biofilm CM were collected on day 1, 3 and 6 of biofilm formation.

### Cell culture and stimulation of macrophages

The murine macrophage cell line RAW 264.7 (ATCC TIB-71, USA) was used for the experiments [26]. Cells were cultivated in DMEM high glucose + 10% heat-inactivated FCS + 1% Penicillin /Streptomycin (Pen/Strep) at 37°C and 5% CO_2_. Cells were transferred into suitable well plate formats at appropriate concentrations (gene expression analysis: 1.5*10^5^ cells / 24 well or 3*10^5^ cells / 12 well; Western Blot: 4*10^5^ cells / 24 well or 8*10^5^ cells / 12 well) and stimulated with the CM 1:1 diluted in fresh cell growth medium. For STING or cGAS inhibition, the cells were pre-incubated with the respective antagonist as suggested by the manufacturer’s protocol (STING: H-151 for 1-2 hours and cGAS: RU.521 for 3 hours, both InvivoGen, USA) in cell growth medium. SA planktonic CM was then added in the same volume for the respective time points. For STING or cGAS activation, the respective agonists (STING: 3’3’-cGAMP and cGAS: G3-YSD/LyoVec^TM^, all InvivoGen, USA) were added to the stimulation with SA biofilm CM. Activation of the IRF3 and NF-κB pathways was evaluated after 4 hours, cytokine induction and IFN-β signaling was analyzed after 20 hours of stimulation. Effect of the inhibitors on cell viability was determined after the maximum incubation period used in the experiments.

IRF3 activation by CM was further validated in primary cells. Therefore, bone marrow cells (BMCs) were isolated from adult C57BL/6 mice in accordance with the national guidelines for animal care. Bone marrow was centrifuged out of the femora and tibiae at 8000 rpm for 3 min and cultivated in α-MEM (Anprotec, Germany) + 10% heat-inactivated FCS + 1% Pen/Strep at 37°C and 5% CO_2_. The next day, suspension cells were harvested, centrifuged and resuspended in respective growth medium. Human peripheral blood mononuclear cells (PBMCs) were isolated from whole blood donated by healthy volunteers upon informed consent via Percoll® density gradient centrifugation, washed and resuspended in RPMI (Anprotec, Germany) + 10% heat-inactivated FCS + 1% Pen/Strep. For both cell types, 10 million cells / 6 well were stimulated with SA planktonic or biofilm CM 1:1 diluted in fresh growth medium (3 ml in total) for 4 hours at 37°C and 5% CO_2_.

### Gene expression analysis of mammalian cells

Total RNA extraction was performed using the innuPREP RNA Mini Kit 2.0 (AJ Innuscreen GmbH, Germany) according to the manufacture’s protocol. Cells were scraped in lysis buffer and transferred to a DNA elimination column. RNA in the lysate was precipitated by adding 70% ethanol, transferred to an RNA column, washed and eluted in H_2_O. Total RNA concentration was measured using the NanoDrop® ND-1000 spectrophotometer (Thermo Scientific, Germany). 1 µg of total RNA was subjected to cDNA synthesis using the RevertUP^TM^ II cDNA synthesis Kit (Biotechrabbit GmbH, Germany) according to the manufacturer’s protocol using Oligo (dT) and hexamer primers. A noRT sample (w/o Reverse Transcriptase) consisting of pooled total RNA of all samples of one experiment was prepared. cDNA was diluted 1:1 in H_2_O and stored at -20°C. 2 µl cDNA template and 400 nM of respective primer pairs were used in qPCR. mRNA levels were evaluated in a two-step PCR reaction (StepOnePlus Real-Time PCR Cycler, Applied Biosystems, USA) with 60°C annealing/extension temperature for 40 cycles using the 2x qPCRBIO SyGreen Mix Hi-ROX (PCR Biosystems Ltd., UK). Quality of qPCR runs and specificity of qPCR products were controlled by included noRT and water samples for each experiment and primer pair and melting curve comparison. mRNA levels of the respective genes of interest (Table 1) were normalized to the reference gene *Hprt1* and calculated by the 2^-ΔCq^ method.

**Table 1.**
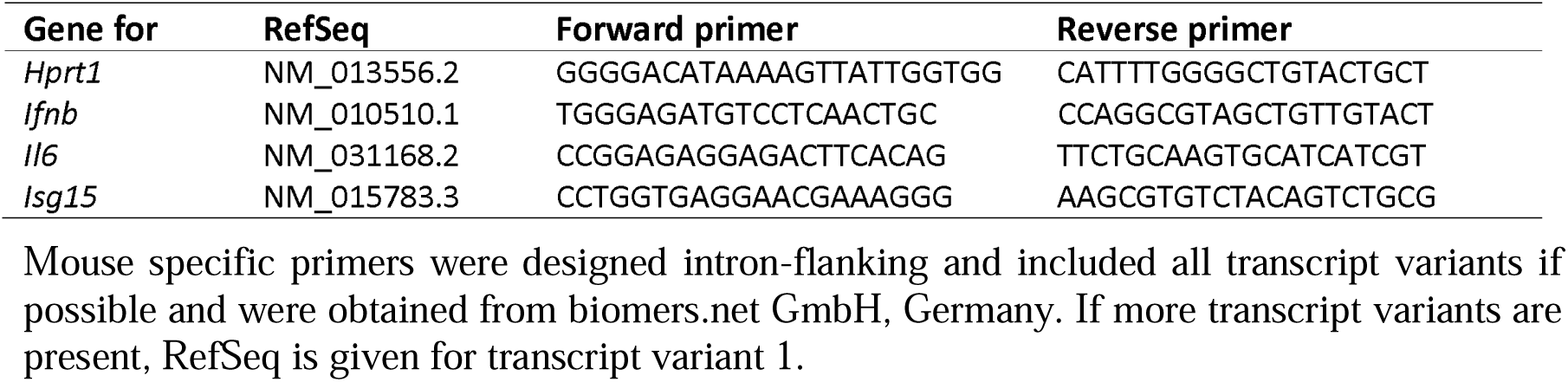
List of mouse specific oligonucleotides used for quantitative RT-PCR analysis.

### Gene expression analysis of bacteria

Planktonic bacteria were harvested by centrifugation of the liquid culture. Biofilms were harvested after removing the supernatants by scraping the biofilm matrix off the well bottom and centrifugation of the pooled material. Pellets were resuspended in TE buffer and treated by sonication (SONOREX SUPER RK 102 H ultrasonic bath, BANDELIN electronic GmbH & Co. KG, Germany) and glass bead disruption (MINI-BEADBEATER™, Bio Spec Products Inc., USA). Lysis was performed with the innuPREP Bacteria Lysis Booster (AJ Innuscreen GmbH, Germany) and subsequent RNA isolation according to the manufacturer’s protocol. RNA samples were additionally treated with DNase I (Promega GmbH, Germany) to eliminate remaining DNA contamination. RNA was transcribed into cDNA by using hexamer primers following the manufacturer’s instructions. qPCR was performed as described above for mammalian cells. mRNA levels of the respective genes of interest (Table 2) were normalized to the reference gene *gyrB* and calculated by the 2^-ΔCq^ method.

**Table 2.**
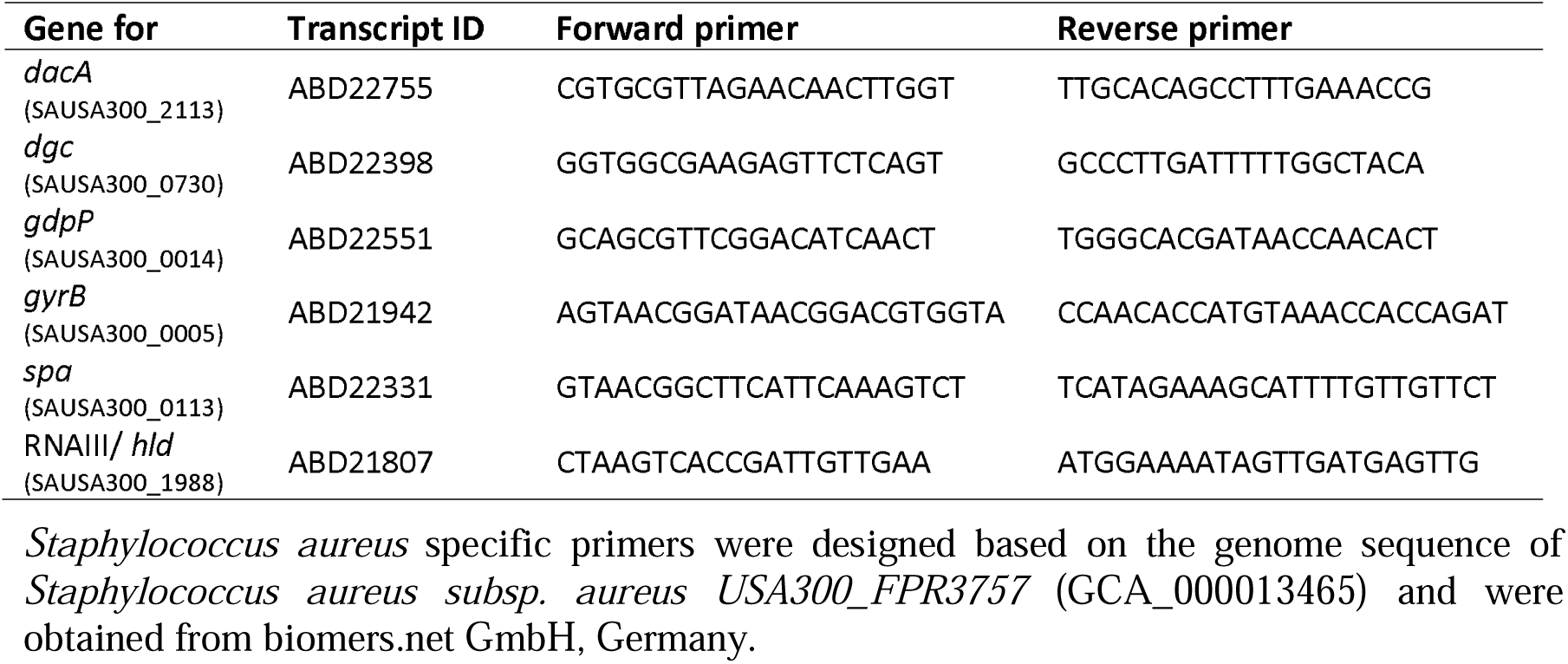
List of bacteria specific oligonucleotides used for quantitative RT-PCR analysis.

**Table 3.** List of antibodies used for Immunoblotting (Western Blot).

| Protein | Source | Size (kDa) | Dilution | Company |
| --- | --- | --- | --- | --- |
| β-Actin | rabbit | 42 | 1:1000 | Proteintech, USA |
| HSP90 | rabbit | 90 | 1:1000 | Cell Signaling Technology, USA |
| Phospho-IRF3 (Ser396) | rabbit | 45-55 | 1:1000 | Cell Signaling Technology, USA |
| Phospho-STAT1 (Tyr701) | rabbit | 84, 91 | 1:1000 | Cell Signaling Technology, USA |
| Anti-rabbit IgG, HRP-linked | goat |  | 1:1000 | Cell Signaling Technology, USA |

Antibodies were all recommended for use in mouse and human (HSP90 and p-IRF3) and applied according to the manufacturer's advice. Proteins were detected by chemiluminescent luminol reaction after incubation with respective HRP-linked secondary antibody and imaged in a ChemoStar ECL Imager.

### Bacterial primer design

SA specific primers were designed based on the genome sequence of *Staphylococcus aureus subsp. aureus USA300_FPR3757* (GCA_000013465). Genome sequences were extracted from the Ensembl Bacteria genome browser (http://bacteria.ensembl.org/index.html) and primers were obtained from biomers.net GmbH, Germany. Primers for the gene for c-di-GMP synthase (diguanylate cyclase, DGC) were designed on the transcript sequence of gene *SAUSA300_0730* which encodes for a protein containing a GGDEF-domain relevant for DGC activity. This gene shows high sequence similarity to gene *SA0701* which already has been associated with c-di-GMP synthesis [27]. Furthermore, the amino acid sequence of SAUSA300_0730 was blasted against the *Staphylococcus aureus* protein database using BLAST®, blastp suite (https://blast.ncbi.nlm.nih.gov/Blast.cgi) and revealed 100% sequence similarity to SA DGC.

### Live/dead staining

After incubation with inhibitors, adherent cells were scraped within the culture medium without prior washing to include dead cells. Cells were centrifuged, washed and resuspended in PBS. 30 nM SYTOX^TM^ Green Nucleic Acid Stain (Invitrogen, USA) was added to the cell suspension and Fluorescein isothiocyanate (FITC)-positive to FITC-negative cell percentage was measured after 5 min of incubation using the BD FACSCanto™ Flow Cytometer (BD Biosciences, USA). Results were analyzed using the Flowing Software (version 2.5.1, Turku Bioscience, Finland).

### Cytometric bead array

Supernatants of gene expression analysis were used for cytometric bead array (CBA, LEGENDplex^TM^, BioLegend, USA) according to the manufacturer’s protocol. A Mouse Inflammation Panel (Mix and Match Subpanel) was used including TNF-α, IL-10, IFN-β and IL-6 beads. In short, supernatants were centrifuged, diluted 1:2.5 with Assay Buffer and together with the standard samples transferred into a V-bottom plate. Bead mix was prepared, added to the samples and incubated on a shaker over night at 4°C in the dark. Plates were then washed twice and incubated with the detection antibody for 1 hour at RT while shaking. Streptavidin-phycoerythrin (SA-PE) was added and further incubated for 30 min at RT while shaking. Plates were washed twice before resuspending the bead pellets in wash buffer. Data acquisition was done with the MACSQuant 10 (Miltenyi Biotech, Germany). Analysis and calculation of cytokine concentrations were performed with the LEGENDplex^TM^ Data Analysis Software (https://legendplex.qognit.com).

### Immunoblotting

Cells were lysed in RIPA buffer (1% v/v NP-40 (IGEPAL® CA-630), 0.25% sodium deoxycholate, 50 mM Tris pH 8.0, 150 mM NaCl, 1 mM EDTA pH 8.0, 1 mM Na_3_VO_4_, 1 mM NaF) including protease inhibitors (Protease Inhibitor Mix G) and phosphatase inhibitors (Phosphatase Inhibitor Mix II, both Serva Electrophoresis GmbH, Germany) for 1 hour at 4°C under rotation. Lysates were centrifuged at 14000 rpm for 20 min at 4°C and protein concentration of supernatants was determined by BCA assay (Cyanagen Srl, Italy10 µg protein was loaded on pre-cast gradient 4-20% Tris-glycine gels (anamed Elektrophorese GmbH, Germany) covered with Laemmli buffer for SDS PAGE (Serva Electrophoresis GmbH, Germany) and separated at 120 V. 5 µl of broad range, color prestained protein standard (New England Biolabs GmbH, Germany) was included on each gel. Proteins were transferred onto an Amersham^TM^ Protran^TM^ 0.45 µm nitrocellulose membrane (GE Healthcare, UK) covered in 4+4 Whatman papers (GE Healthcare, UK) soaked in transfer buffer (192 mM glycine, 25 mM Tris, 2.6 mM SDS, 0.5 mM NA_3_VO_4_, 15% v/v methanol) for 1 hour at 2 mA/cm^2^. After Ponceau S stain (0.5 % Ponceau S, 3 % trichloroacetic acid, 96.5 % ddH_2_O), membranes were cut according to sizes of investigated proteins to allow detection of multiple proteins. Membranes were blocked with BlueBlock PF (Serva Electrophoresis GmbH, Germany) for 30 min at RT with continuous shaking. Incubation with primary antibodies (listed in Table 2) was performed in BlueBlock PF over night at 4°C. After three times washing with TBST, membranes were incubated with the respective HRP-linked secondary antibody (Table 2) for 1 hour at RT under shaking and again washed three times. Blots were developed with ECL substrate (WESTAR ETA C ULTRA 2.0, Cyanagen Srl, Italy) and imaged in the ChemoStar ECL & Fluorescence Imager (Intas Science Imaging Instruments GmbH, Germany).

### Quantification of Protein A in CM

Protein A was quantified in the CM using the Protein A ELISA Kit (abcam, UK) according to the manufacturer’s protocol. In short, CM were diluted 1:50 for Medium and biofilm CM and 1:5000 for planktonic CM. Samples were pretreated with a boiling step in sample diluent + enhancer (Kit components) for 5 minutes and standard was prepared. Samples and standard were transferred into the ready-to-use ELISA plate. Antibody cocktail was prepared and added to the wells. The plate was incubated for 1 h at RT. After 3 washing steps, TMB development solution was added and incubated for 10 min. Stop solution was added and OD at 450 nm was detected using the CLARIOstar® Plus (BMG Labtech, Germany).

### Quantification of 3’3’-cGAMP in CM

3’3’-cGAMP was quantified in the CM using the DetectX® 3’3’-cGAMP ELISA Kit (Arbor Assays, USA) according to the manufacturer’s protocol. In short, CM were used undiluted and standard was prepared. Samples and standard were transferred into the ready-to-use ELISA plate. DetectX® 3’,3’-cGAMP Conjugate and Antibody were added to the wells. The plate was incubated for 1 h at RT with shaking. After 4 washing steps, TMB substrate was added and incubated for 30 min. Stop solution was added and OD at 450 nm was detected using the CLARIOstar® Plus (BMG Labtech, Germany).

### Statistical analysis

Experiments were done in n = 3 - 6 independent replicates as stated in the figure legends. Data are presented as mean ± SEM and single values as dots. Statistical evaluation was performed using ordinary one-way ANOVA with post-hoc multiple comparison testing and Bonferroni correction. A p-value below 0.05 was considered statistically significant. Data analysis was performed with GraphPad Prism for Windows (Version 9.3.1, GraphPad Software Inc., USA).

## Results

### Inhibition of STING prevents IRF3 mediated *Ifnb* induction by macrophages upon stimulation with SA planktonic CM

Stimulation of RAW 264.7 macrophages with CM of SA planktonic or biofilm cultures showed that the SA planktonic environment activates the IRF3 pathway whereas p-IRF3 was not detected in the respective biofilm environment (Fig. 1A upper blot). This effect was also observed in primary mouse BMCs and human PBMCs (Fig. 1A middle and lower blot) indicating that IRF3 pathway activation upon stimulation with SA planktonic CM is conserved over different cell lines and species. As shown in Figure 1B-E, STING inhibition by the small molecule inhibitor H-151 dose-dependently prevented induction of *Ifnb* and its target gene *Isg15*, in line with the absence of IRF3 and STAT1 activation upon stimulation with SA planktonic CM. *Il6* mRNA levels were only slightly and non-significantly reduced by H-151 (Fig. 1F) indicating that the general inflammatory response independent of IFN-β was still induced by the SA planktonic environment. H-151 treatment showed adverse effects on cell viability at high concentrations (Fig. 1G). However, the used range of 4 to 400 ng/ml had no (4 and 40 ng/ml) or still tolerable (400 ng/ml) effects on cell viability. Thus, our data indicate that STING activation is crucial for the IRF3 induced IFN-β response in a SA planktonic environment.

**Fig. 1.**
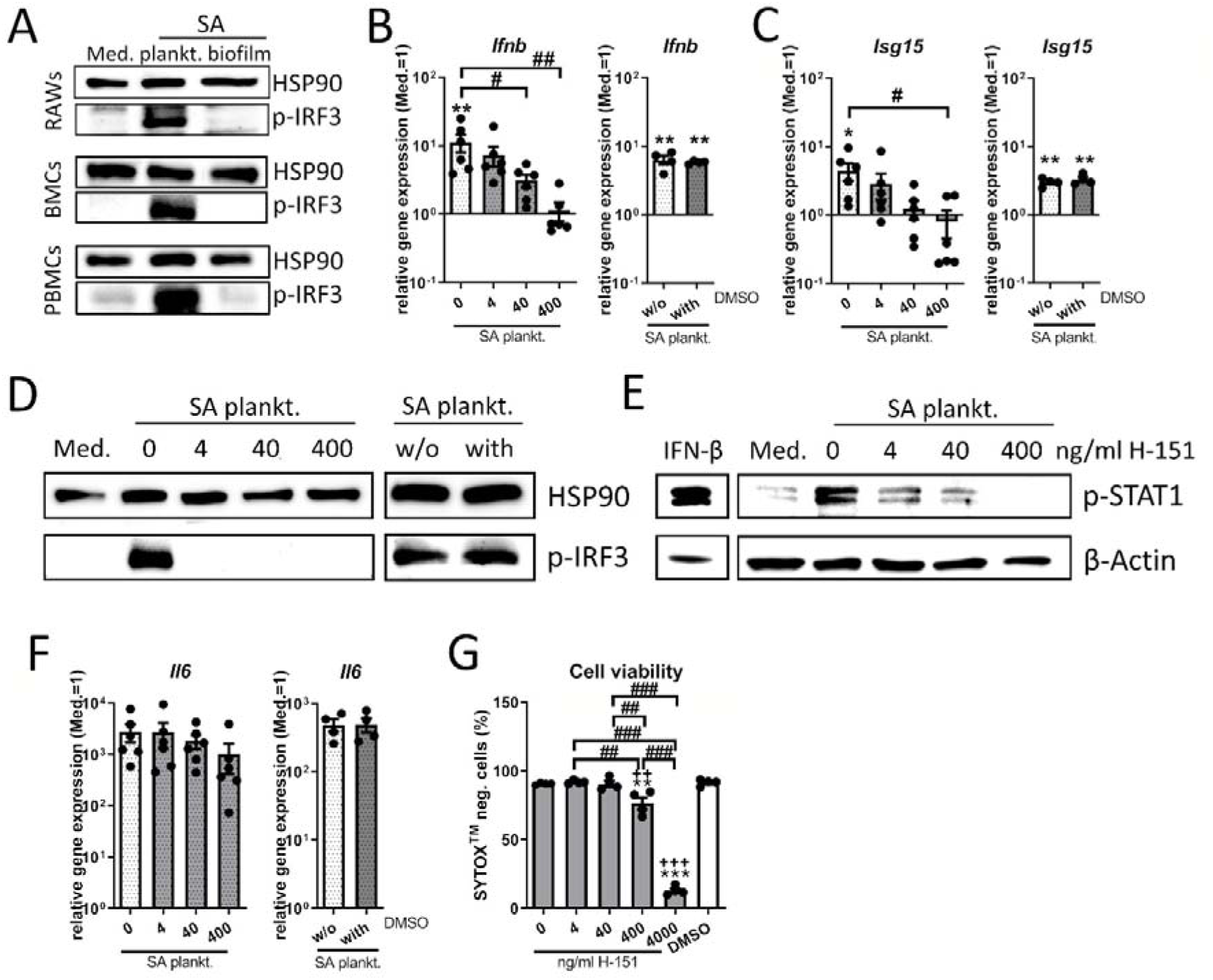
**Effect of STING inhibition on the IRF3/IFN-**β **response upon stimulation with SA planktonic CM.** Cells were stimulated with SA planktonic CM 1:1 diluted in fresh growth media (DMEM high glucose + 10% FCS + 1% Pen/Strep) and effect of the STING antagonist H-151 on the IRF3 mediated IFN-β response was evaluated. A) Activation of the IRF3 pathway in different cell populations. RAW 264.7 cells, primary mouse BMCs and primary human PBMCs were treated with SA planktonic or biofilm CM for 4 hours and presence of phospho-IRF3 as activated form of the transcription factor was visualized by Western Blot. HSP90 was used as loading control. n=3 experiments. B+C) Gene expression analysis of *Ifnb* and target gene *Isg15*. RAW 264.7 cells were treated with H-151 in different concentrations (4, 40 and 400 ng/ml) and SA planktonic CM for 20 hours and mRNA levels of *Ifnb* (B) and *Isg15* (C) were quantified by RT-qPCR. DMSO as solvent control was added to SA planktonic CM in the volume according to the highest inhibitor concentration. Data are presented as relative gene expression of gene of interest related to the reference gene *Hprt1* and normalized to the unstimulated medium control. n=4 (DMSO) or 6 (H-151) experiments. D) Activation of the IRF3 pathway. RAW 264.7 cells were treated with H-151 in different concentrations (4, 40 and 400 ng/ml) and SA planktonic CM for 4 hours and presence of phospho-IRF3 as activated form of the transcription factor was visualized by Western Blot. DMSO as solvent control was added to SA planktonic CM in the volume according to the highest inhibitor concentration. HSP90 was used as loading control. n=3 experiments. E) Activation of the IFN-β pathway. RAW 264.7 cells were treated with H-151 in different concentrations (4, 40 and 400 ng/ml) and SA planktonic CM for 20 hours and presence of phospho-STAT1 as activated form of the transcription factor and downstream target of the IFN-β receptor was visualized by Western Blot. β-Actin was used as loading control. n=3 experiments. F) Gene expression analysis of stress marker *Il6*. RAW 264.7 cells were treated with H-151 in different concentrations (4, 40 and 400 ng/ml) and SA planktonic CM for 20 hours and mRNA levels of *Il6* were quantified by RT-qPCR. DMSO as solvent control was added to SA planktonic CM in the volume according to the highest inhibitor concentration. Data are presented as relative gene expression of gene of interest related to the reference gene *Hprt1* and normalized to the unstimulated medium control. n=4 (DMSO) or 6 (H-151) experiments. G) Viability of cells after treatment with H-151. RAW 264.7 cells were treated with H-151 in different concentrations (4, 40, 400 and 4000 ng/ml) for 22 hours (duration of the experiments covering 2 hours of pre-incubation and 20 hours of CM stimulation) and cell viability was measured by SYTOX^TM^-staining and FACS analysis. Percentage of SYTOX^TM^- negative (living) cells are shown. n=4 experiments. For B+C/F+G: Data are presented as mean ± SEM and single values are shown as dots. p-values were calculated by ordinary one- way ANOVA with post-hoc Bonferroni corrected multiple comparison. * is indicating significance against Medium, + is indicating significance against DMSO, # is showing significance between inhibitor concentrations. * p<0.05, ** p<0.01, *** p<0.001; + p<0.05, ++ p<0.01, +++ p<0.001; # p<0.05, ## p<0.01, ### p<0.001.

### STING activation leads to an IRF3 induced IFN-**β** response by macrophages upon stimulation with SA biofilm CM

We next wanted to investigate if the SA biofilm environment actively inhibited STING activation and if that was the cause of the missing IRF3 activation. Therefore, we added the bacterial STING agonist 3’3’-cGAMP to the stimulation with SA biofilm CM and evaluated IRF3 activation and the subsequent induction of an IFN-β response. 3’3’-cGAMP significantly induced *Ifnb* and *Isg15* gene expression upon stimulation with SA biofilm CM (Fig. 2A). In line with this, IRF3 was also found to be phosphorylated in the samples that had been treated with 3’3’-cGAMP in the presence of SA biofilm CM (Fig. 2B). *Il6* gene expression was induced by SA biofilm CM and further increased when 3’3’-cGAMP was added (Fig. 2C). The observed IFN-β induction and increased IL-6 expression by 3’3’- cGAMP were corroborated on the protein levels (Fig. 2D). Protein levels of the pro- inflammatory cytokine TNF-α were not affected, whereas protein levels of the anti- inflammatory cytokine IL-10 were significantly elevated by 3’3’-cGAMP addition to SA biofilm CM resulting in a significant decrease of pro-inflammatory macrophage polarization (Fig. 2F). Interestingly, the effects of 3’3’-cGAMP were significantly higher in combination with SA biofilm CM than alone. This indicates that the SA biofilm environment is not preventing STING pathway activation, and that addition of 3’3’-cGAMP to the SA biofilm CM could lead to a more pronounced IFN-β response.

**Fig. 2.**
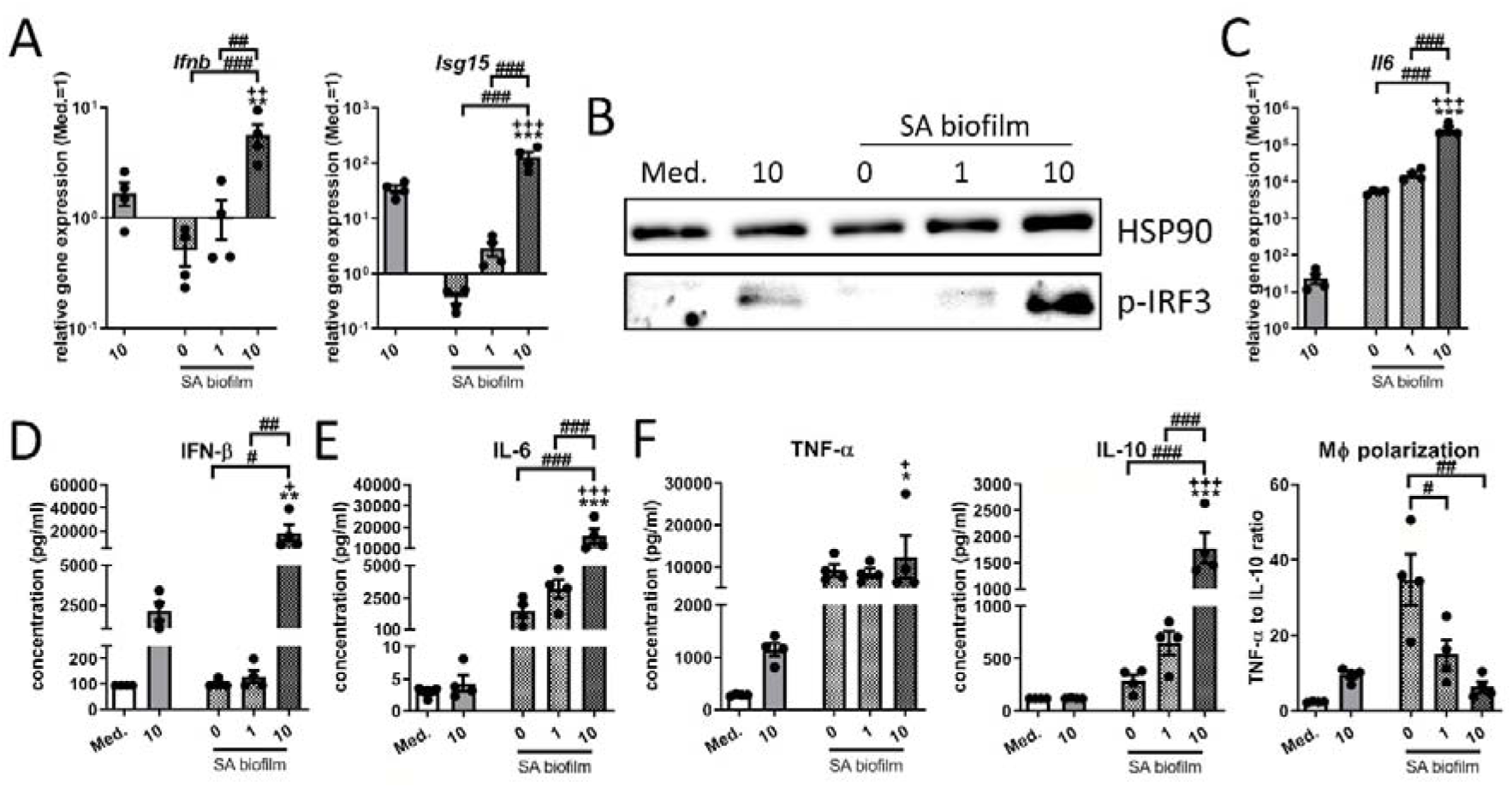
**Effect of STING activation on the IRF3/IFN-**β **response upon stimulation with SA biofilm CM.** RAW 264.7 cells were stimulated with SA biofilm CM 1:1 diluted in fresh growth media (DMEM high glucose + 10% FCS + 1% Pen/Strep) and effect of the bacterial STING agonist 3’3’-cGAMP on the IRF3 mediated IFN-β response was evaluated. A) Gene expression analysis of *Ifnb* and target gene *Isg15*. RAW 264.7 cells were treated with 3’3’- cGAMP in different concentrations (1 and 10 µg/ml) and SA biofilm CM for 20 hours and mRNA levels of *Ifnb* (B) and *Isg15* (C) were quantified by RT-qPCR. Data are presented as relative gene expression of gene of interest related to the reference gene *Hprt1* and normalized to the unstimulated medium control. n=4 experiments. B) Activation of the IRF3 pathway. RAW 264.7 cells were treated with 3’3’-cGAMP in different concentrations (1 and 10 µg/ml) and SA biofilm CM for 4 hours and presence of phospho-IRF3 as activated form of the transcription factor was visualized by Western Blot. HSP90 was used as loading control. n=3 experiments. C) Gene expression analysis of stress marker *Il6*. RAW 264.7 cells were treated with 3’3’-cGAMP in different concentrations (1 and 10 µg/ml) and SA biofilm CM for 20 hours and mRNA levels of *Il6* were quantified by RT-qPCR. Data are presented as relative gene expression of gene of interest related to the reference gene *Hprt1* and normalized to the unstimulated medium control. n=4 experiments. D-E) Cytokine release of IFN-β, IL-6, pro- inflammatory TNF-α and anti-inflammatory IL-10. RAW 264.7 cells were treated with 3’3’- cGAMP in different concentrations (1 and 10 µg/ml) and SA biofilm CM for 20 hours and protein concentrations of IFN-β (D) and IL-6 (E) as well as pro-inflammatory TNF-α and anti-inflammatory IL-10 (E) were quantified in the supernatant by cytometric bead array (CBA; LEGENDplex^TM^). Data are presented as absolute concentration (pg/ml). Ratio of TNF- α to IL-10 protein levels was used as indicator for macrophage polarization. n=4 experiments. For A/C-F: Data are presented as mean ± SEM and single values are shown as dots. p-values were calculated by ordinary one-way ANOVA with post-hoc Bonferroni corrected multiple comparison. * is indicating significance against Medium, + is indicating significance between 10 µg/ml 3’3’-cGAMP in Medium or SA biofilm CM, # is showing significance between 3’3’-cGAMP concentrations. * p<0.05, ** p<0.01, *** p<0.001; + p<0.05, ++ p<0.01, +++ p<0.001; # p<0.05, ## p<0.01, ### p<0.001.

### Gene expression levels associated with virulence but not CDN synthesis are decreasing with biofilm formation and *Ifnb* induction

Our data indicated that the stimulation with SA planktonic CM leads to STING activation whereas the required stimulus seemed to be absent in the respective biofilm CM. Bacteria- derived CDNs are known ligands for STING [9] and also play a role in bacterial adaption processes such as biofilm formation [11]. Thus, we evaluated the effect of biofilm formation on the gene expression levels of CDN synthases at different time points during biofilm formation. As shown in Figure 3A, both, planktonic and biofilm SA expressed the genes encoding the CDN synthases for c-di-AMP (diadenylate cyclase *dacA*) and c-di-GMP (diguanylate cyclase *dgc*) as well as the phosphodiesterase *gdpP*, the enzyme that degrades c- di-AMP. The mRNA levels of *dacA* did not differ significantly between planktonic and biofilm SA, whereas the gene expression levels of diguanylate cyclase *dgc* were highest in day 1 biofilm SA. In line, the mRNA levels of the phosphodiesterase *gdpP* were lowest in day 1 biofilm SA and increased with biofilm age (Fig. 3A). We could not detect relevant amounts of the bacterial CDN 3’3’-cGAMP in the SA CM (Fig. 3B). These data indicate that CDNs are not the main mediators of STING/IRF3 activation and *Ifnb* induction in SA planktonic CM. We further investigated gene expression levels of molecules associated with QS and virulence as these are known to be regulated in the process of SA biofilm formation [28, 29]. Here, we found that mRNA levels of the effector molecule of the accessory global regulator (agr) QS system RNAIII, and the RNAIII encoded gene of the phenol soluble modulin δ-toxin (*hld*) significantly decreased during biofilm formation (Fig. 3C). Gene expression levels of *sarS*, a member of the staphylococcal accessory regulator (SarA) QS system and direct activator of Protein A (*spa*) [30] did not differ significantly between planktonic and biofilm cultures but was lowest in day 1 biofilm SA (Fig. 3D). Expression of Protein A significantly decreased with biofilm formation and maturation on mRNA and protein level (Fig. 3E). Overall, gene expression analysis of planktonic and biofilm bacteria revealed that mRNA levels of CDN synthases did not correlate with the IRF3 mediated *Ifnb* induction in SA planktonic CM. Instead, our data show that genes associated with QS and virulence were highly expressed in planktonic SA and down-regulated with biofilm formation suggesting that bacterial factors other than CDNs might be involved in the STING pathway activation.

**Fig. 3.**
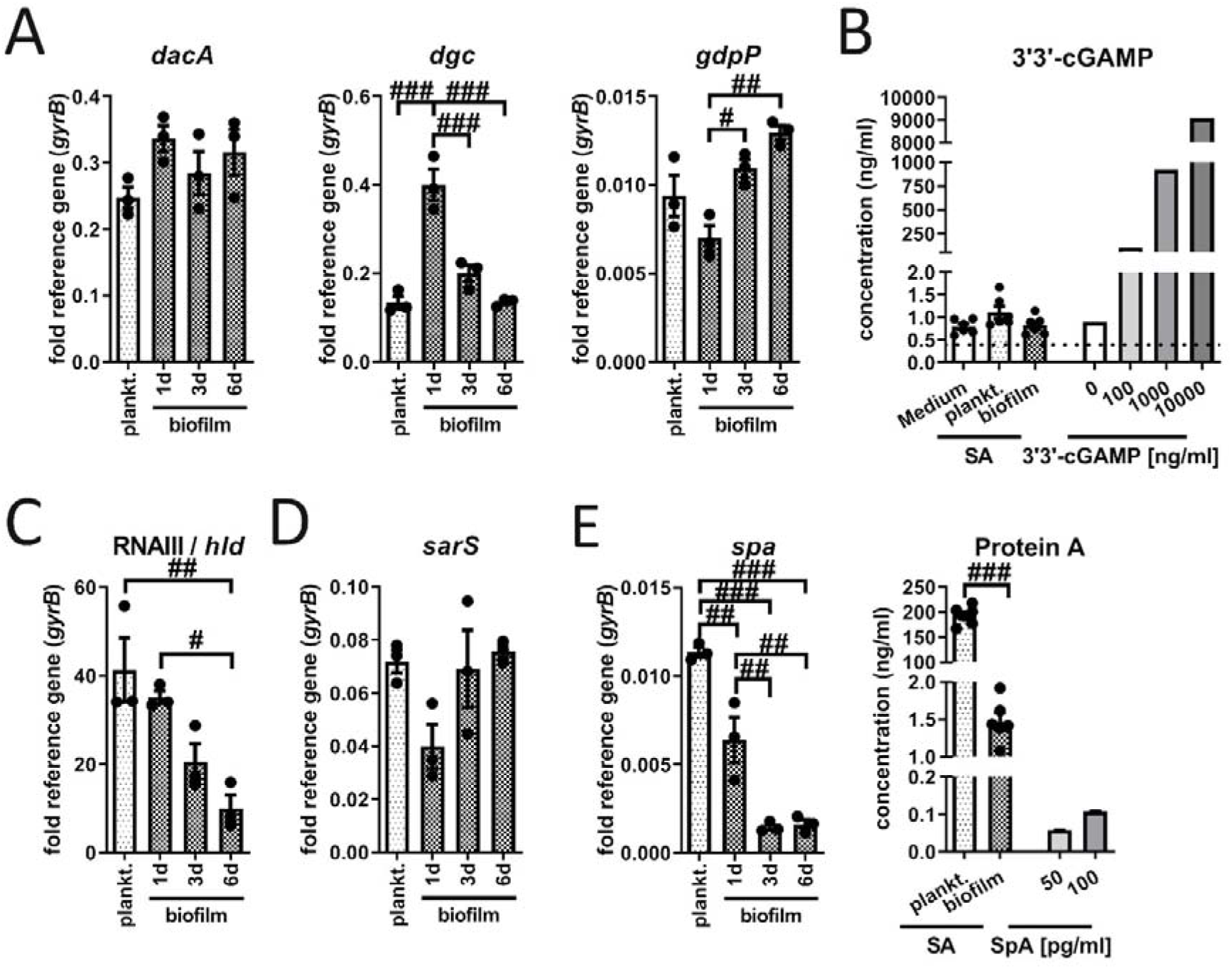
Differences in bacterial gene expression and release of 3’3’-cGAMP or Protein A between SA planktonic and biofilm cultures. SA was cultured in DMEM high glucose with 10% heat-inactivated FCS at 37°C and 5% CO_2_ either under shaking for planktonic or static for biofilm conditions. Bacteria and supernatants were harvested after 24 hours of planktonic or after 1 day, 3 days or 6 days of biofilm culture. A) Gene expression analysis of c-di-AMP synthase (diadenylate cyclase, DacA), c-di-GMP synthase (GGDEF domain protein, diguanylate cyclase, DGC) and phosphodiesterase (GdpP). mRNA levels of *dacA*, *dgc* and *gdpP* were quantified by RT-qPCR. Data are presented as relative gene expression of gene of interest related to the reference gene *gyrB*. n=3 experiments. B) 3’3’-cGAMP release by planktonic and biofilm SA. 3’3’-cGAMP concentration was measured in Medium without bacteria, SA planktonic CM and 6 days biofilm CM. For validation of the assay, known concentrations of 3’3’-cGAMP (0, 100, 1000 and 10000 ng/ml) in medium were included in the measurement. Concentrations were quantified in the supernatants by competitive ELISA. Data are presented as absolute concentration (ng/ml). n=3 different CM batches in duplicates. C+D) Gene expression analysis of agr / SarA quorum sensing (QS) molecules. mRNA levels of the agr effector molecule RNAIII and RNAIII encoded PSM δ-toxin (*hld* gene, C) and SarA regulator protein *sarS* (D) were quantified by RT-qPCR. Data are presented as relative gene expression of gene of interest related to the reference gene *gyrB*. n=3 experiments. E) Protein A production by planktonic and biofilm SA. mRNA levels of *spa* were quantified by RT-qPCR. Data are presented as relative gene expression of gene of interest related to the reference gene *gyrB*. n=3 experiments. Protein A concentration was measured in SA planktonic CM and 6 days biofilm CM. For validation of the assay, known concentrations of Protein A (SpA; 50 and 100 pg/ml) in medium were included in the measurement. Concentrations were quantified in the supernatants by quantitative Sandwich ELISA. Data are presented as absolute concentration (ng/ml). n=3 different CM batches in duplicates. For all: Data are presented as mean ± SEM and single values are shown as dots. p-values were calculated by ordinary one-way ANOVA with post-hoc Bonferroni corrected multiple comparison. # is showing significance between different CM. # p<0.05, ## p<0.01, ### p<0.001.

### Cytosolic DNA sensor cGAS is involved in the IRF3 mediated *Ifnb* induction upon stimulation with SA planktonic CM

As direct STING activation by bacteria-derived CDNs seemed not to be the mechanism behind the IRF3 mediated *Ifnb* induction in SA planktonic CM, we investigated if the cytosolic DNA sensor cGAS, upstream of STING, was involved. Therefore, we inhibited cGAS by treating the cells with the small molecule inhibitor RU.521 prior to CM stimulation. Figure 4A shows that inhibition of cGAS by RU.521 efficiently and completely blocked the activation of the cGAS pathway by its agonist G3-YSD. Cell viability was not affected by the treatment with the cGAS inhibitor RU.521 at the used concentrations (Fig. 4B). Furthermore, cGAS inhibition resulted in a dose-dependent reduction of *Ifnb* and *Isg15* induction upon stimulation with SA planktonic CM (Fig. 4C). Induction of *Il6* gene expression levels were only slightly but not significantly reduced by cGAS inhibition (Fig. 4D). Unexpectedly, adding the cGAS agonist G3-YSD to the stimulation with SA biofilm CM was not sufficient to induce gene expression of *Ifnb* and its target gene *Isg15* compared to cGAS activation by G3-YSD in the medium control (Fig. 4E). Induction of *Il6* gene expression by SA biofilm CM was not affected by the cGAS agonist (Fig. 4F). Thus, unlike for direct STING activation with 3’3’-cGAMP, the treatment of the cells with the cGAS agonist G3-YSD did not result in induction of *Ifnb* gene expression and downstream targets upon stimulation with SA biofilm CM. This suggests that the necessary stimulus is not only missing in the biofilm environment, but even that the biofilm environment prevents cGAS activation.

**Fig. 4.**
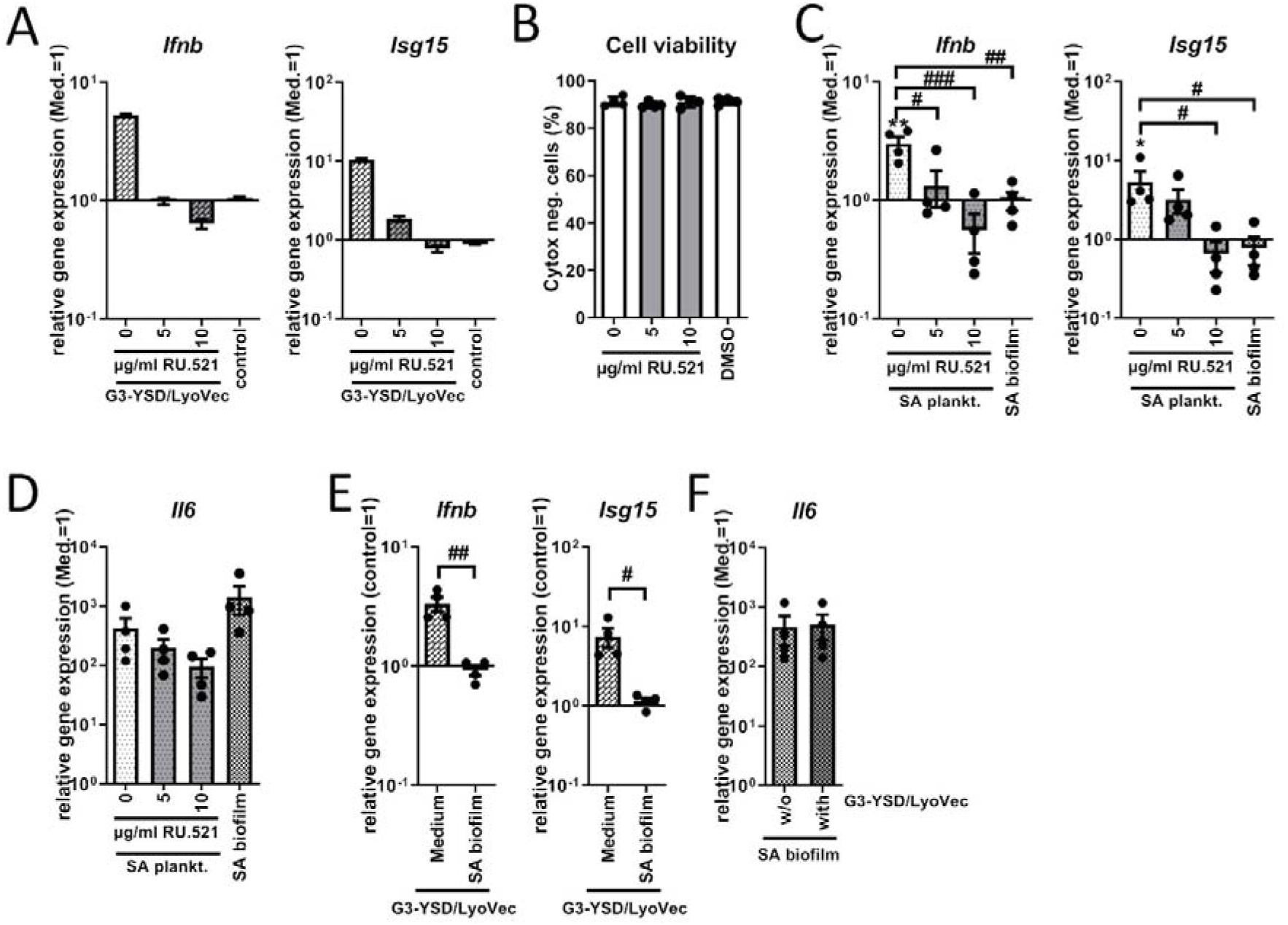
Role of cGAS on the *Ifnb* induction upon stimulation with SA planktonic or biofilm CM. Cells were stimulated with SA planktonic or biofilm CM 1:1 diluted in fresh growth media (DMEM high glucose + 10% FCS + 1% Pen/Strep) and effect of the cGAS antagonist RU.521 or cGAS agonist G3-YSD/LyoVec^TM^ on the induction of *Ifnb* and its target gene expression was evaluated. A) Evaluation of the cGAS inhibitor RU.521. RAW 264.7 cells were treated with RU.521 in different concentrations (5 and 10 µg/ml) and 100 ng/ml of the cGAS agonist G3-YSD or its control was added to the cells together with the transfection reagent LyoVec^TM^ for 20 hours. mRNA levels of *Ifnb* and *Isg15* were quantified by RT-qPCR. Data are presented as relative gene expression of gene of interest related to the reference gene *Hprt1* and normalized to the unstimulated medium control. n=2 experiments. B) Viability of cells after treatment with RU.521. RAW 264.7 cells were treated with RU.521 in different concentrations (5 and 10 µg/ml) for 23 hours (duration of the experiments covering 3 hours of pre-incubation and 20 hours of CM stimulation) and cell viability was measured by SYTOX^TM^ -staining and FACS analysis. DMSO as solvent control was added in the volume according to the highest inhibitor concentration. Percentage of SYTOX^TM^ - negative (living) cells are shown. n=4 experiments. C+D) Gene expression analysis of *Ifnb*, target gene *Isg15* and stress marker *Il6*. RAW 264.7 cells were treated with RU.521 in different concentrations (5 and 10 µg/ml) and SA planktonic CM was added to the cells for 20 hours. Cells stimulated with SA biofilm CM were included for comparison. mRNA levels of *Ifnb* and *Isg15* (C) and *Il6* (D) were quantified by RT-qPCR. Data are presented as relative gene expression of gene of interest related to the reference gene *Hprt1* and normalized to the unstimulated medium control. n=4 experiments. E+F) Gene expression analysis of *Ifnb*, target gene *Isg15* and stress marker *Il6*. RAW 264.7 cells were treated with medium or SA biofilm CM and 100 ng/ml of the cGAS agonist G3-YSD or its control together with the transfection reagent LyoVec^TM^ for 20 hours. mRNA levels of *Ifnb* and *Isg15* (E) and *Il6* (F) were quantified by RT-qPCR. Data are presented as relative gene expression of gene of interest related to the reference gene *Hprt1* and normalized either to the control/LyoVec^TM^ sample (*Ifnb* + *Isg15*) or to the unstimulated medium control (*Il6*). n=4 experiments. For B-F: Data are presented as mean ± SEM and single values are shown as dots. p-values were calculated by ordinary one-way ANOVA with post-hoc Bonferroni corrected multiple comparison. * is indicating significance against Medium, # is showing significance between different inhibitor concentrations or treatments. * p<0.05, ** p<0.01, *** p<0.001; # p<0.05, ## p<0.01, ### p<0.001.

### Induction of *Ifnb* gene expression by SA planktonic CM is not conserved throughout different SA strains

To understand if the observed effects were dependent on the characteristics of the specific SA strain used for our CM preparation, we next included MRSA strain USA300 and reference SA strain ATCC 25923 in our study. Therefore, we cultivated the different strains side by side and in the same conditions and prepared CM from planktonic and biofilm cultures of different ages. Interestingly, the induction of *Ifnb* and *Isg15* gene expression which we had observed for planktonic CM using the strain ATCC 42930 directly disappeared as early as day 1 when changing the culture conditions from planktonic to biofilm. For CM generated from USA300, the highest induction of *Ifnb* and *Isg15* gene expression was detected in planktonic CM but was still detectable on day 1 with biofilm CM and then declined during biofilm maturation until day 6. Stimulation with both, planktonic or biofilm CM of the SA strain ATCC 25923 and also SA strain SA113 (data not shown) did not result in induction of *Ifnb* and *Isg15* gene expression at all (Fig. 5A). Cells responded to all CM with increased *Il6* mRNA levels, indicating that planktonic as well as the biofilm environments of all SA strain were able to induce an adequate immune response (Fig. 5B). *Ifnb* induction by planktonic CM of USA300 was dependent on STING activation (Fig. 5C) whereas induction of *Il6* gene expression was not affected by STING inhibition (Fig. 5D) which is in line with the results received for the SA strain ATCC 49230. USA300 (FPR3757) was isolated from a wrist abscess and ATCC 49230 (UAMS-1) from an osteomyelitis and therefore, both can be considered as invasive and virulent SA strains. Taken together, these results indicate that the STING dependent *Ifnb* induction is likely associated with the degree of virulence and the release of bacterial factors into the environment.

**Fig. 5.**
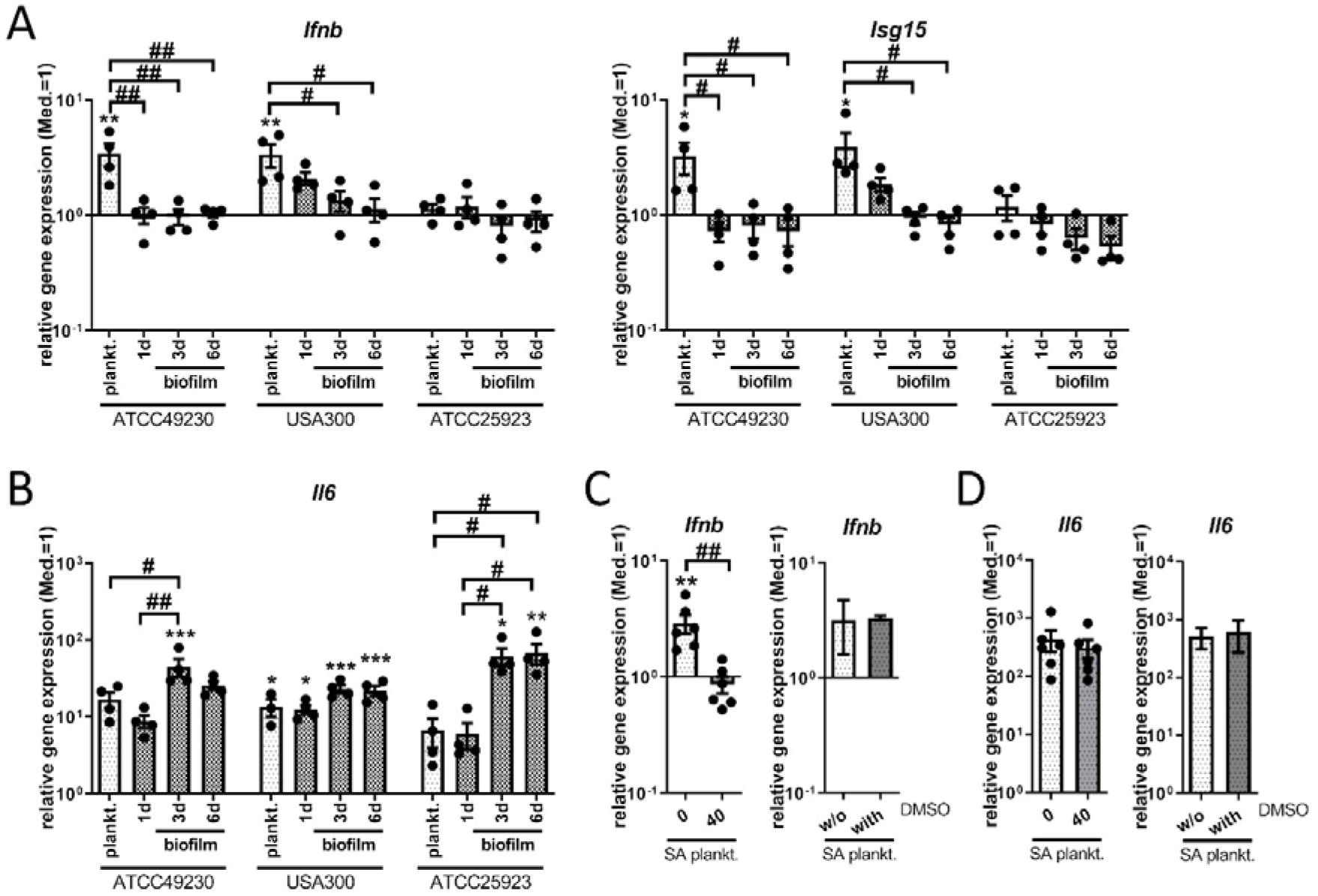
*Ifnb* induction upon stimulation with SA planktonic or biofilm CM of different SA strains. CM were generated from planktonic or biofilm cultures of SA strains ATCC 49230, the MRSA USA300 and the reference strain ATCC 25923. Planktonic CM were harvested after 24 hours of shaking culture and biofilm CM were harvested on day 1 (young), day 3 (maturating) and day 6 (mature) of static culture. RAW 265.7 cells were stimulated with CM 1:1 diluted in fresh growth media (DMEM high glucose + 10% FCS + 1% Pen/Strep) and induction of an IFN-β response was evaluated. A+B) Gene expression analysis of *Ifnb*, target gene *Isg15* and stress marker *Il6*. RAW 264.7 cells were stimulated with the CM for 20 hours and mRNA levels of *Ifnb* and *Isg15* (A) and *Il6* (B) were quantified by RT-qPCR. C+D) STING inhibition on gene expression analysis of *Ifnb*, target gene *Isg15* and stress marker *Il6*. RAW 264.7 cells were treated with 40 ng/ml H-151 and SA planktonic CM generated from the MRSA strain USA300 for 20 hours and mRNA levels of *Il6* (D) were quantified by RT- qPCR. DMSO as solvent control was added to SA planktonic CM in the volume according to the inhibitor concentration. Data are presented as relative gene expression of the gene of interest related to the reference gene *Hprt1* and normalized to the unstimulated medium control. n=2 (DMSO) or 6 (H-151) experiments. For all: Data are presented as mean ± SEM and single values are shown as dots. p-values were calculated by ordinary one-way ANOVA with post-hoc Bonferroni corrected multiple comparison. * is indicating significance against Medium, # is showing significance between planktonic and biofilm CM or inhibitor concentrations. * p<0.05, ** p<0.01, *** p<0.001; + p<0.05, ++ p<0.01, +++ p<0.001; # p<0.05, ## p<0.01, ### p<0.001.

## Discussion

An impaired immune response during biofilm formation is a major problem of implant-related bone infections and an attractive target for innovative therapeutic strategies. In a previous study, we found that the presence of an SA planktonic environment induces IFN-β production, presumably via IRF3 activation, which was not detectable in the respective biofilm environment [6]. As STING activation leads to an IRF3 mediated production of IFN-β and the STING/IRF3 pathway gains increasing attention in bacterial infection research, we investigated if STING was involved in the observed *Ifnb* induction upon stimulation with SA planktonic CM. Indeed, we found that the *Ifnb* induction by SA planktonic CM was dependent on STING and that STING activation resulted in *Ifnb* gene expression in SA biofilm CM. This could not be explained by different expression levels of CDN synthases by planktonic or biofilm SA. Furthermore, cGAS seemed to play a role in the *Ifnb* induction by SA planktonic CM, but addition of a cGAS agonist in contrast to direct activation of STING was not sufficient to induce *Ifnb* expression in biofilm CM. Instead, our results indicated that the expression levels of genes associated with virulence were involved. Interestingly, *Ifnb* gene expression was detected upon stimulation with planktonic CM generated from the two virulent and invasive SA strains ATCC 49230 and USA300, whereas this was not observed for the SA strains ATCC 25923 and SA113. Overall, our data indicate that STING activation is associated with a higher degree of virulence. As virulence decreases during conversion to the biofilm lifestyle, increasing biofilm maturation prevents *Ifnb* induction.

Our data are in line with other studies showing a STING dependent IFN-β production in RAW 264.7 cells or mouse BMDMs upon infection with clinically relevant SA strains [23, 31]. STING activation was only observed for living SA and disappeared when bacteria were heat-killed indicating that the stimulus had to be actively produced by the bacteria. Hence, it can be assumed that in our indirect infection model the relevant bacterial factor is produced during the planktonic culture and released into the environment. C-di-NMPs are discussed to play a role in the initiation of the switch from a motile to a sessile lifestyle of bacteria and by this might be involved in the immune recognition of early biofilm formation via activation of the STING pathway [32]. Indeed, c-di-AMP released by SA biofilms was already shown to induce an IFN-β response in macrophages [24]. Thus, we investigated if differences in the c- di-NMP production between planktonic and biofilm SA could be the relevant factor in our model system. In line with the other study, the diadenylate cyclase *dacA* was found to be expressed in our SA biofilm cultures. Furthermore, we found mRNA levels of the diguanylate cyclase *dgc* to be increased in day 1 biofilm SA which fits to the function of c-di-GMP as a factor involved in planktonic to biofilm transition [32, 33]. Controversially, the gene expression levels of CDN synthases that we observed did not correlate with the ability to cause an *Ifnb* induction. In addition, we did not detect *Ifnb* gene expression upon stimulation with biofilm CM indicating that bacterial CDNs might not been the mechanism behind the STING dependent *Ifnb* induction observed in our model.

Biofilm formation is further associated with a change in gene expression towards environmental adaption and reduced virulence [5, 34]. Thus, we compared expression levels of the agr QS effector molecule RNAIII, the encoded δ-toxin (*hld*) and Protein A (*spa*) between planktonic and biofilm SA. In line with the existing literature, expression of these virulence associated genes was highest in planktonic SA and decreased with progressing biofilm formation. Additionally, high c-di-GMP concentrations are associated with reduced expression of virulence factors [35]. As *dgc* mRNA levels were highest during early biofilm SA, this further indicated that the virulence of biofilm cultured SA is decreased. Thus, it seems that the observed STING dependent *Ifnb* induction might be related to the different (?) production of bacterial factors associated with virulence. Protein A for example was able to induce IFN-β production in airway epithelial cells upon endocytic uptake [36]. Additionally, the ability to induce an IFN-β response via the NOD2-IRF5 pathway was associated with the virulence of the respective SA strain [37]. To strengthen our theory that a different release of bacterial factors associated with virulence is the main mediator resulting in *Ifnb* induction via the STING/IRF3 pathway in a planktonic but not in the biofilm environment, we aimed to exclude an inhibitory effect of the biofilm CM. In a previous study, we had already shown that low glucose and high lactate levels characteristic for the biofilm environment did not prevent IRF3 activation and *Ifnb* induction of SA planktonic CM [6]. Furthermore, addition of the bacterial STING agonist 3’3’-cGAMP resulted in a pronounced IRF3 mediated IFN-β production in SA biofilm CM. This indicates that the stimulus for STING activation is missing in the SA biofilm environment.

As our results suggested that bacterial CDNs are not responsible for a direct STING activation in the SA planktonic environment, we investigated the role of the upstream cytosolic DNA sensor cGAS. cGAS was already described to be involved in STING activation upon SA infections [23, 38]. Indeed, the inhibition of cGAS resulted in the loss of *Ifnb* induction upon stimulation with SA planktonic CM. cGAS can either be activated by foreign DNA derived from invading microbes or by endogenous DNA released from damaged and dying host cells and then lead to subsequent STING activation via synthesis of 2’3’-cGAMP [16, 39]. In a previous study, we showed that SA biofilm CM contained more free bacterial DNA than the respective planktonic CM [40] indicating that bacterial DNA in the environment cannot be the cause for cGAS activation by SA planktonic CM. Furthermore, uptake of surrounding bacterial DNA normally requires endocytosis and results in activation of TLR-9 signaling [41, 42]. Thus, it is rather likely that increased production of virulence factors by planktonic SA induces host cell damage and release of endogenous DNA. This is in line with our previous finding that stimulation with planktonic CM of the less virulent *Staphylococcus epidermidis* (SE) did not result in IRF3 activation [6] and that *Ifnb* induction is dependent on the virulence of the SA strain. Additionally, the study by Scumpia et al. showed that IFN-β production via the cGAS/STING pathway increased with increasing virulence of the staphylococci strains which the authors also explained with a higher rate of cell death [23]. Unexpectedly, addition of the cGAS agonist G3-YSD resulted in *Ifnb* gene expression in medium but was not sufficient to induce *Ifnb* gene expression in SA biofilm CM. So far, we cannot explain exactly how the biofilm environment prevents cGAS activation and to clarify this is part of ongoing investigations.

The influence of IFN-β on the outcome of the immune response is controversially discussed for bacterial infections [43, 44]. In SA pneumonia, IFN-β is associated with suppressed immune cell recruitment and cytokine production resulting in increased morbidity [36, 37]. In a SA skin infection model, however, IFN-β supported an effective immune response and enhanced bacterial clearance [45]. Thus, it has to be further clarified if the activation of the cGAS/STING/IRF3 pathway and the subsequent production of IFN-β is of a beneficial or detrimental nature. Besides IFN-β production, STING activation further results in an increased production of IL-6 and IL-10 upon stimulation with SA biofilm CM. It was shown that the anti-inflammatory cytokine IL-10 contributes to bacterial persistence in a SA orthopedic biofilm infection model [46]. Thus, induction of an IFN-β response in a SA biofilm situation via STING activation has to be carefully evaluated and the possibility that anti-inflammatory processes and immune tolerance are supported need to be taken into consideration.

Overall, our results show that the cGAS/STING/IRF3 pathway is differentially regulated in SA planktonic and biofilm environments and therefore might be an attractive target for immunomodulatory interventions that try to strengthen the immune response in the course of biofilm formation. Nevertheless, the role of IFN-β in this process has to be fully clarified to assess whether STING inhibition or activation is a conceivable approach also in the treatment of implant-related bone infections.

## Author contribution

ES was responsible for study conception and design, acquisition, analysis and interpretation of data and wrote the manuscript. GS participated in data acquisition and analysis. KFK contributed to data interpretation and critically revised the manuscript. All authors read and approved the final manuscript.

## Acknowledgements

We would like to thank Dafina Kadrijaj and Tabea Elschner for their help with experiments. Furthermore, we thank Dr. Lan-Sun Chen for providing human PBMCs.

## Data availability statement

The data that support the findings of this study are available from the corresponding author upon reasonable request.

